# Resolvin D2/GPR18 signaling enhances monocytic myeloid-derived suppressor cell function to mitigate abdominal aortic aneurysm formation

**DOI:** 10.1101/2024.02.23.581672

**Authors:** Paolo Bellotti, Zachary Ladd, Victoria Leroy, Gang Su, Shiven Sharma, Joseph B. Hartman, Jonathan Krebs, Chelsea Viscardi, Robert Maile, Lyle L. Moldawer, Phillip Efron, Ashish K. Sharma, Gilbert R. Upchurch

## Abstract

Abdominal aortic aneurysm (AAA) formation is a chronic vascular pathology characterized by inflammation, leukocyte infiltration and vascular remodeling. The aim of this study was to delineate the protective role of Resolvin D2 (RvD2), a bioactive isoform of specialized proresolving lipid mediators, via G-protein coupled receptor 18 (GPR18) receptor signaling in attenuating AAAs. Importantly, RvD2 and GPR18 levels were significantly decreased in aortic tissue of AAA patients compared with controls. Furthermore, using an established murine model of AAA in C57BL/6 (WT) mice, we observed that treatment with RvD2 significantly attenuated aortic diameter, pro-inflammatory cytokine production, immune cell infiltration (neutrophils and macrophages), elastic fiber disruption and increased smooth muscle cell α-actin expression as well as increased TGF-β2 and IL-10 expressions compared to untreated mice. Moreover, the RvD2-mediated protection from vascular remodeling and AAA formation was blocked when mice were previously treated with siRNA for GPR18 signifying the importance of RvD2/GPR18 signaling in vascular inflammation. Mechanistically, RvD2-mediated protection significantly enhanced infiltration and activation of monocytic myeloid-derived suppressor cells (M-MDSCs) by increasing TGF-β2 and IL-10 secretions that mitigated smooth muscle cell activation in a GPR18-dependent manner to attenuate aortic inflammation and vascular remodeling via this intercellular crosstalk. Collectively, this study demonstrates RvD2 treatment induces an expansion of myeloid-lineage committed progenitors, such as M-MDSCs, and activates GPR18-dependent signaling to enhance TGF-β2 and IL-10 secretion that contributes to resolution of aortic inflammation and remodeling during AAA formation.

## INTRODUCTION

Abdominal aortic aneurysms (AAA) are a devastating clinical disease and a leading cause of mortality, especially among the elderly population in the US, with no specific medical therapies available(1–4). The pathological and immunological process of AAAs is characterized by a multi-faceted crosstalk between leukocyte infiltration into the aortic wall, such as neutrophils and macrophages, with subsequent destruction of elastin and collagen in the medial and adventitial layers of the aorta due to excessive production of pro-inflammatory cytokines and matrix degrading enzymes(1, 5–7). This chronic non-resolving inflammatory process during aging, termed as ‘inflammaging’, culminates in aortic smooth muscle cell (SMC) activation and remodeling of the aorta that can result in aortic wall thinning with eventual rupture and sudden death(8, 9).

The resolution of aortic inflammation is an active process that is dependent on the balance of pro-inflammatory (leukotrienes, prostaglandins, cytokines) and pro-resolving (specialized proresolving lipid mediators; SPMs) paracrine mediators(10). An important bioactive isoform of SPMs is Resolvin D2 (RvD2) which is derived from omega-3 fatty acids, such as docosahexaenoic acid (DHA)(11). RvD2 promotes a pro-resolving and pro-reparative phenotype via G-protein coupled receptor 18 (GPR18) to regulate phagocytosis and cytokine production(12). Our previous study demonstrated that treatment with RvD2 significantly influences macrophage polarization towards an anti-inflammatory M2 phenotype to prevent AAA formation(13). However, the role of monocytes to reprogram towards a more pro-resolving and pro-reparative phenotype in chronic vascular inflammation remains to be deciphered. Furthermore, the cell-specific mechanism of RvD2-mediated resolution via GPR18 signaling also remains to be investigated in the pathogenesis of AAAs.

Recent evidence suggests that myeloid-derived suppressor cells (MDSCs) are an immunosuppressive cell population that are generated during acute and chronic pathologies secondary to infections, sterile inflammation or tumor biology (14–16). This heterogenous cell population, comprising of monocytic-(M-) and granulocytic (G- or PMN-)-MDSCs, are generated by enhanced myelopoiesis and characterized by myriad of immune modulating activities, including release of cytokines, exhaustion of pro-inflammatory cells, and promotion of pro-resolving cell phenotypes (17–20). These immature progenitor cells are notably distinguished for their ability to act in an immunosuppressive manner, ultimately modulating the survival, function, and proliferation of other immune cells(18, 21–31). Therefore, the enhancement of endogenous immunosuppression via M-MDSCs represents a significant signaling axis that could potentially be exacerbated by pharmacological modalities.

In this study, the immunomodulation of vascular inflammation and remodeling during AAA formation by RvD2 and GPR18 signaling was investigated using an established topical elastase-treatment model of experimental murine AAA. Aortic wall from human patients undergoing AAA repair demonstrated a marked decrease in RvD2 expression. In addition, treatment with RvD2 in the experimental murine models of AAA showed preserved aortic morphology and reduced inflammatory cytokine expression. RvD2 treatment correlated with increased M-MDSCs in the aortic tissue and *in vitro* studies demonstrated the ability of this monocytic subset to downregulate aortic smooth muscle cell activation to attenuate vascular remodeling.

## MATERIALS AND METHODS

### Animals

Eight to 12-week old wild-type C57BL/6 male mice (Jackson Laboratory, Bar Harbor, ME) were housed and maintained at 70°F, 50% humidity, in 12-hour light-dark cycles as per institutional animal protocols. Mice were provided drinking water and standard chow diet *ad libitum*. All experiments were approved by and conducted in accordance to the Institutional Animal Care and Use Committee of the University of Florida (protocol # 201910902).

### Human aortic tissue analysis

Collection of human aortic tissue was approved by the University of Florida’s Institutional Review Board (#IRB201902782). Consent was obtained from all patients before surgery. Aortic tissue from AAA patients was resected during open surgical repair as well as during organ transplant donor surgery (controls). Aortic tissue was homogenized in Trizol, and RNA was purified per manufacturer’s protocol (Qiagen, Valencia, CA). cDNA was synthesized using the iScript cDNA Synthesis Kit (BioRad, Hercules, CA). Quantitative Real-Time (RT-PCR) was performed with primer sets (MWG/Operon, Huntsville, AL) in conjunction with SsoFast EvaGreen Supermix (BioRad, Hercules, CA). Gene expression was calculated by using the relative quantification method according to the following equation: 2(−ΔCT), where ΔCT=(Average gene of interest)−(Average reference gene), where GAPDH was used as the reference gene.

### Topical Elastase AAA Model

C57BL/6 (wild-type; WT) mice were anesthetized using isoflurane and underwent exposure of the infrarenal abdominal aorta, as previously described(32). The aorta was dissected circumferentially away from the surrounding tissues and subjected topically for 5 minutes to either 5 µL of elastase (0.4 U/mL type 1 porcine pancreatic elastase, Sigma Aldrich, St. Louis, MO) or heat-inactivated elastase (deactivated elastase was used as control). Treated aortic sections were harvested postoperatively and preserved in formalin for immunohistochemistry or snap-frozen in liquid nitrogen for protein and RNA extraction. Aortic diameter was measured using video micrometry (AmScope, Irvine, CA). Aortic dilation percentage was determined by [(maximal AAA diameter – self-control aortic diameter)/(self-control aortic diameter)] × 100. An aortic dilation of ≥100% was considered positive for AAA.

### Treatment of Mice with RvD2

To evaluate the attenuative effects of RvD2 in AAA formation, mice received intraperitoneal injections (i.p.) with 0.3 mL vehicle (0.1% ethanol in 0.9% saline) or RvD2 (300ng/kg; Cayman Chemical, Ann Arbor, MI) on postoperative days 1 through 13 and subsequently aortas were harvested on post-operative day 14.

### GPR18 Expression Quantification

To quantify expression of the GPR18 receptors, RNA was isolated from the aortic samples using TRIzol Reagent (Ambion, Carlsbad, CA). cDNA was synthesized from the isolated RNA via the iScript Reverse Transcription supermix (BioRad, Hercules, CA) and quantitative (real-time) RT-PCR was performed with primer sets (MWG/Operon, Huntsville, AL) in conjunction with SsoFast EvaGreen Supermix (BioRad, Hercules, CA), as previously described(33). To evaluate expression of GPR18 receptors, the following primers were used for evaluating human aortic tissue: GPR18, Forward: GCCAAGCGTTACACTGGAAA; Reverse: TGATACTTAGAAACTCCTGTCCATC, and the following primers were used for murine aortic tissue: GPR18 Forward: TCATGATCGGGTGCTACGTG; GPR18 Reverse: CTTGTAGCATCAGGACGGCA, GAPDH Forward: TTGATGGCAACAATCTCCAC; GAPDH Reverse: CGTCCCGTAGACAAAATGGT. Gene expression was calculated by using the relative quantification method according to the following equation: 2(−ΔCT), where ΔCT = (average gene of interest) − (average reference gene), where GAPDH was used as the reference gene. Each PCR reaction was carried out in triplicate, and the relative quantification of gene expression was quantified as fold change.

### *In vivo* Suppression of GPR18

Mice were injected i.p. with siRNA targeting murine GPR18 (10nmol; ON-TARGETplus SMARTPool siRNA, Dharmacon, Lafayette, CO) or a non-targeting control siRNA (5nmol; ON-TARGETplus Non-targeting Control siRNA, Dharmacon, Lafayette, CO). The day after injection, mice underwent surgery using the topical elastase model to induce AAAs, as previously described(34). Aortas were harvested on day 14 and expression of GPR18 was measured by RT-qPCR from isolated RNA, as described above. RvD2 was injected i.p. from days 1 to 13 postoperatively in mice that received siRNA for GPR18 and non-targeting controls. Aortic tissue was harvested on postoperative day 14 for further analysis and aortic dilation percentage was measured.

### Histology

Harvested aortic tissue was fixed in 10% zinc-buffered formalin overnight, processed with ethanol, and embedded into paraffin. Slides of aortic cross-sections were prepared and stained for elastin (Van Gieson’s Solution, catalog no. s289; Poly Scientific R&D Systems, Bay Shore, NY) smooth muscle actin (monoclonal anti-actin α-smooth muscle, Sigma Aldrich, St. Louis, MO), macrophages (purified anti-mouse Mac-2, Cedarlane Laboratories, Burlington, Canada), and neutrophils (rat anti-mouse neutrophils, AbD Serotec, Oxford, UK). Images were taken with 20X magnification by a Nikon microscope equipped with a digital camera and Elements BR software. Staining was quantified by measuring the percent intensity of positive sections relative to the whole aortic cross-section using ImagePro software (Media Cybernetics, Rockville, MD, USA).

### Enzyme-linked Immunosorbent Assay

RvD2 was quantified by ELISA using the supplied manufacturer’s protocol (Cayman Chemical, Ann Arbor, MI).

### Flow Cytometry Analysis

Murine aortic tissue was harvested from WT mice after undergoing AAA induction on days 3, 7 and 14, as previously described(34). Tissue was minced and incubated for 15 min at 37°C with collagenase type IA (Sigma Aldrich; St. Louis, MO) in PBS with 0.5% BSA and 2mM EDTA Following red blood cell lysis, the cell suspension was stained with Live/Dead (Aqua; Biolegend, San Diego, CA) diluted in PBS for 30min at 4°C. Cells were then washed in PBS and incubated with FC block (BD Biosciences) for 5 min followed by an extracellular antibody staining cocktail for 30min. Cells were washed twice and then fixed with fixation buffer (eBioscience; Thermo Fisher Scientific) for 25 min at 4°C. Following fixation, cells were washed in permeabilization buffer (eBioscience) and incubated with intracellular iNOS antibody for 30 min at 4°C. Flow cytometry was performed on a BD FACSymhony™A3 Cell Analyzer (BD Biosciences) and data was analyzed using FlowJo Software. M-MDSCs were identified as CD45^+^CD11b^+^CD11c^-^Ly6C^+^Ly6G^-^iNOS^+^ and G-MDSCs were identified as CD45^+^CD11b^+^CD11c^-^Ly6C^-^Ly6G^+^iNOS^-^, as previously described (35–37) and quantified as a percentage of total CD45^+^ cells.

### *In vitro* generation of M-MDSCs

M-MDSCs were generated for *in vitro* experiments as previously described (37, 38). Briefly, cells were isolated from bone marrow of C57Bl/6 mice and cultured with granulocyte-macrophage stimulating factor (100ng or 2×10^3^ units per 3×10^6^ cells, Peprotech, Cranbury, NJ) in RPMI-1640 media containing 10% heat inactivated fetal bovine serum, 1% antibiotic-antimycotic, 1% L-glutamine, and 0.1% β-mercaptoethanol for 3 days. The resulting cell culture, consisting of M- and G-MDSCs, was further separated into Ly6G^-^ and Ly6G^+^ subpopulations with Anti-Ly6G UltraPure Microbeads (Miltenyi Biotec) via AutoMACS magnetic based cell sorting. The Ly6G^-^ or Ly6G^+^ population, representing M-MDSCs and G-MDSCs respectively, was further cultured in the presence of LPS (0.1mg/mL; Invitrogen) and IFN-γ (100 U/mL; BioLegend, San Diego, California) for 24hrs and subjected to a Dead Cell Removal kit via magnetic separation (Miltenyi Biotec) and live M- or G-MDSCs were subsequently used for *in vitro* experiments.

### Cell Culture Experiments

C57BL/6 murine M-MDSCs and aortic smooth muscle cells (SMCs) (ATCC, Manassas, VA) were cultured (2×10^5^ cells in 48-well plates) in DMEM F12 with 10% FBS. Cells were transfected for 6 hours with GPR18 siRNA, or a non-targeting control siRNA as described above, in serum-free media containing a transfection reagent (HiPerFect Transfection Reagent, catalog no. 301704; Qiagen, Valencia, CA). M- or G-MDSC cultures were subjected to a transient elastase treatment (0.4 U) for 5 minutes, washed, and treated with media containing either RvD2 (300nM) or a vehicle control. After 24hrs supernatant was collected and the cytokine expression was analyzed by luminex bead array assay (Millipore Sigma, Burlington, MA). In separate cultures, conditioned media transfer (CMT) was performed using M-MDSCs and SMCs. M-MDSCs were treated with or without RvD2 for 6 h, and CMT was performed to SMC cultures treated with/without elastase, and culture supernatants were harvested after 24 h for analysis of cytokines (CXCL1 and IL-6) and MMP2 activity (Luminex bead array, Millipore Sigma, St. Louis, MO).

### Statistical Analysis

Statistical analysis was performed with GraphPad 10 (GraphPad Software, La Jolla, CA). One-way analysis of variance (ANOVA) with Tukey’s multiple comparisons test was used to compare means between three or more groups. An unpaired Student’s t-test with nonparametric Mann-Whitney or Wilcoxon rank sum test was used for pairwise comparison of groups. Results are displayed as mean ± standard error of mean with p<0.05 considered as statistically significant.

## RESULTS

### Resolvin D2 attenuates experimental murine AAA formation

Using the topical elastase model, mice were treated with either elastase or heat-inactivated elastase on day 0 and injected with vehicle or RvD1 (300ng/kg bodyweight) daily on post-operative days 1 to 13 and harvested on day 14 (**Figure 1A**). A significant increase in mean aortic diameter was noted in elastase treated mice compared to heat-inactivated elastase treated control mice (209±4.3% vs 1.2±0.3%; n=10-19/group; p<0.0001; **Figure 1B-C**). When mice were administered RvD2, elastase-treated mice demonstrated significantly decreased mean aortic dilation compared to mice treated with vehicle alone (141.4±5.0 vs. 209±4.3%; n=19-20/group; p< 0.0001; **Figure 1B-C**). Additionally, RvD2 treated mice demonstrated significantly increased expression of smooth muscle alpha actin (SM-αA), as well as decreased expression of elastic breaks and immune cell (neutrophils and macrophages) infiltration on day 14 compared to mice treated with vehicle alone (**Figure 1D-E**). The expression of pro-inflammatory cytokines (IFN-γ, IL-1β, IL-6, TNF-α, IL-17, MCP-1, MIP-2, HMGB1) and matrix metalloproteinase (MMP2) were significantly attenuated by RvD2 treatment (**Figure 2**). Moreover, the expression of TGF-β2 (38.8±7.4 vs. 5.5±1.6 pg/ml; p<0.001) and IL-10 (235±26.8 vs. 98.5±9.6 pg/ml; p<0.001) were significantly increased in the RvD2-treated mice compared to untreated controls (**Figure 2**). Analysis of human aortic tissue demonstrated significantly decreased expression of RvD2 and GPR18 expression in human aortic tissue from AAA patients compared to controls (**Supplemental Figure S1**).

**Figure 1.**
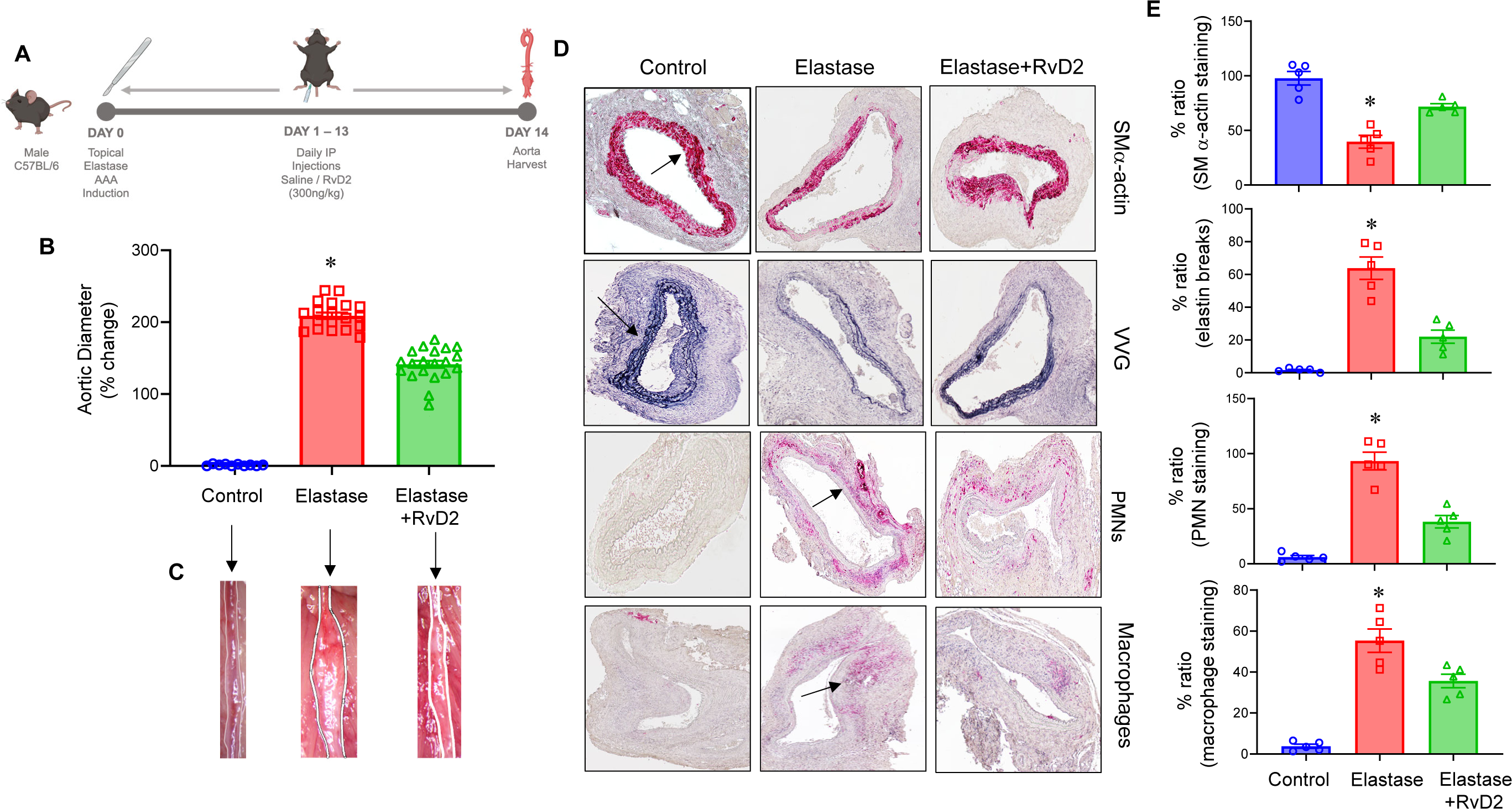
RvD2 administration decreases aortic aneurysm phenotype and preserves morphology to mitigate AAA formation. **A**, Schematic description of murine topical elastase model with RvD2 treatments. Mice were divided into three groups and treated with either heat-inactivated elastase or elastase on day 0. Mice were then administered either vehicle or RvD2. Aortic diameter was measured on day 14 and tissue harvested for additional analysis. **B**, RvD2 treated mice demonstrated a significant decrease in aortic diameter compared to vehicle treated mice; *p<0.001 vs. other groups; n=10-20 per group). **C**, Representative images of aortic phenotype in the respective groups. **D**, Comparative histology performed on day 14 indicates that elastase-treated mice administered with RvD2 have a marked increase in smooth muscle cell α-actin (SM-α actin) expression, decrease in elastic fiber disruption (Verhoeff-Van Gieson staining for elastin) as well as neutrophil (PMN) and macrophage (Mac-2) infiltration, compared to elastase-treated WT mice alone (n=5 per group). Representative histological images in the respective groups with arrows indicate areas of immunostaining. **E**, Quantification of histological staining in respective groups; n=5 per group; *p<0.02 vs. other groups.

**Figure 2.**
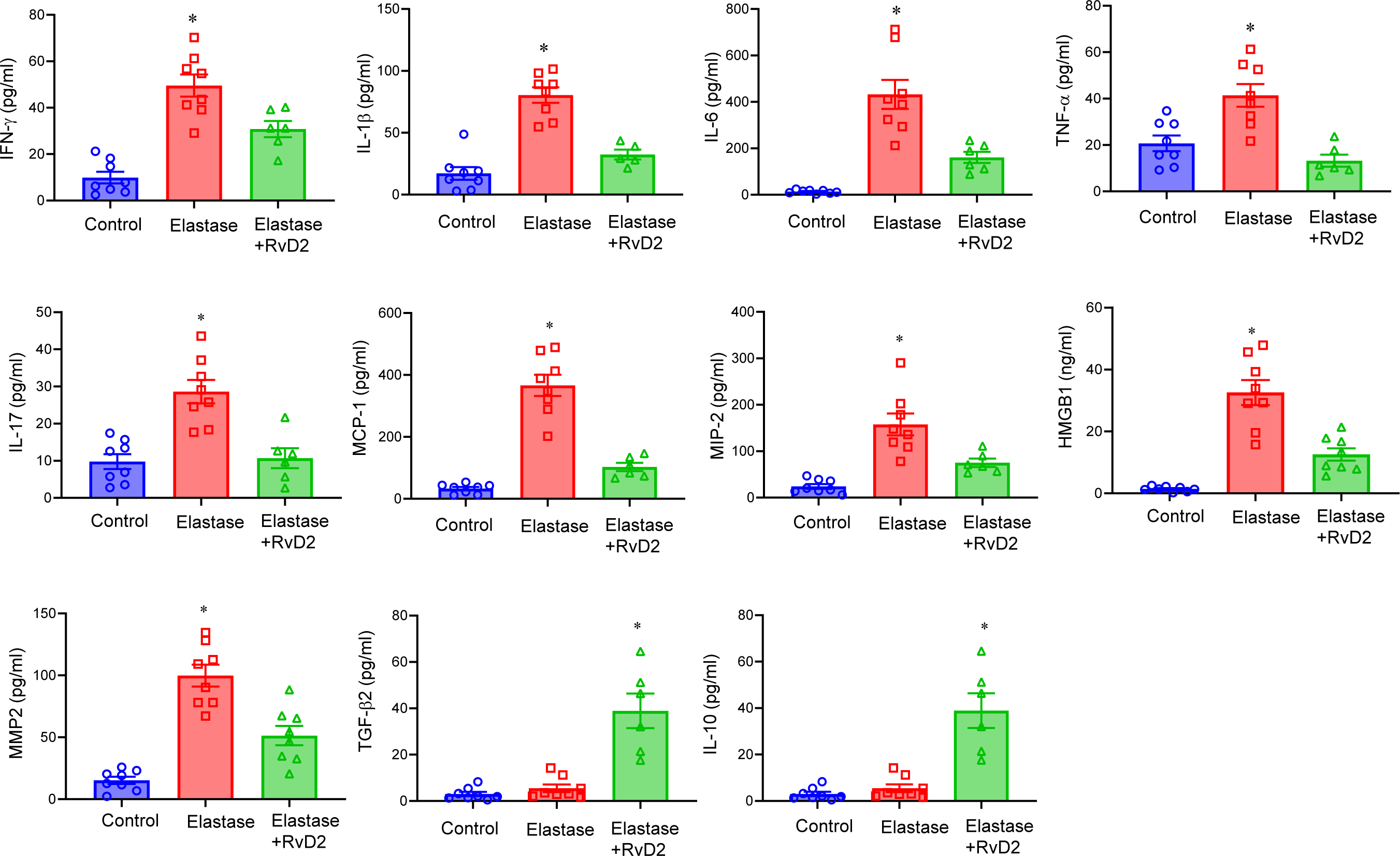
Pro-inflammatory cytokine production is decreased by RvD2 treatment. Aortic tissue from RvD2-treated mice showed a significant attenuation in pro-inflammatory cytokine/chemokine production and MMP2 expression compared to elastase-treated WT mice alone. There was a significant increase in the expression of anti-inflammatory cytokines, TGF-β2 and IL-10, between RvD2-treated mice compared to untreated mice. *p<0.01 vs. other groups; n=6-8 mice per group.

### RvD2-mediated protection is mediated by GPR18-dependent signaling to mitigate AAA formation

In order to assess the molecular target and ligand-receptor interactions of RvD2, the effect of *in vivo* knockdown of endogenous murine GPR18 was examined. Mice injected with GPR18 siRNA had a significantly reduced GPR18 expression compared to non-targeting controls (**Supplemental Figure 2**). Treatment with GPR18 siRNA obliterated the protective effect of RvD2 as observed by increased aortic diameter compared to mice administered with control siRNA+RvD2 (169.5±8.4% vs. 137.2±5.6%; p=0.004, **Figure 3A-B**) Expression of smooth muscle-α actin was significantly increased, while elastin fragmentation, as well as neutrophil and macrophage infiltration were significantly decreased in mice administered with control siRNA+RvD2 compared to GPR18-siRNA+RvD2 treated animals (**Figure 3C-D**). The expression of pro-inflammatory cytokines (IFN-γ, IL-1β, IL-6, TNF-α, IL-17, MCP-1, MIP-2, HMGB1) and matrix metalloproteinase (MMP2) were significantly attenuated, while the expression of anti-inflammatory TGF-β2 (47.4 ±14.9 vs. 10.9±2.0 pg/ml; p<0.01) and IL-10 (223.9±34.7 vs. 29.0±8.1 pg/ml; p=0.007) were significantly increased by RvD2 treatment in control siRNA treated mice but not in GPR18 siRNA-treated mice (**Figure 4**).

**Figure 3.**
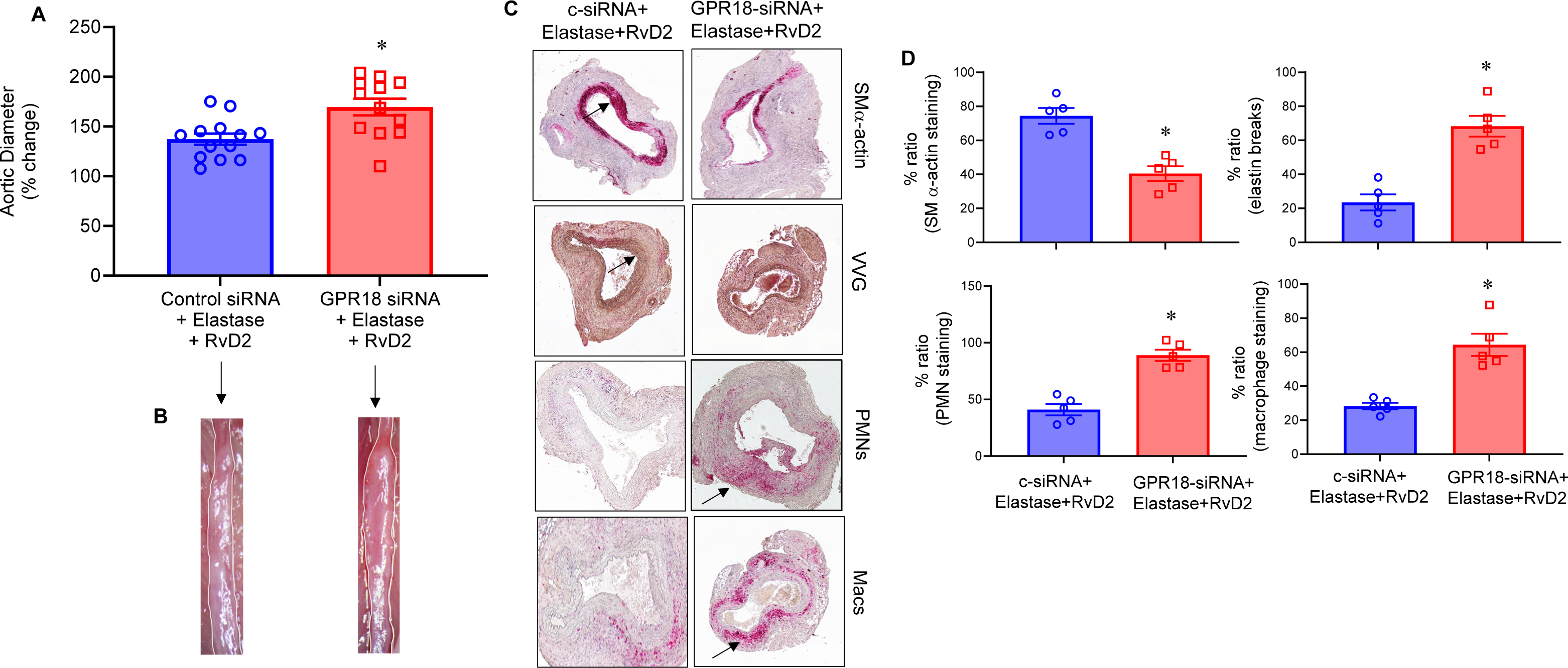
*In vivo* GPR18 knockdown reduces the protective effect of RvD2. **A**, GPR18-siRNA treatment of WT mice demonstrated a significant increase in aortic diameter as compared to control (c)-siRNA treated mice after administration of RvD2 in respective groups (*p=0.004, n=12-13 per group). **B**, Representative images of aortic phenotype in respective groups. **C,** Expression of smooth muscle-α actin is significantly increased, and elastin fragmentation as well as macrophage and neutrophil infiltration in aortic tissue are significantly decreased in mice treated with c-siRNA+RvD2 compared to mice treated with GPR18 siRNA+RvD2 (n=5 per group). Arrows indicate areas of immunostaining. **D,** Quantification of histological staining in respective groups; n=5 per group; *p<0.01 vs. other groups.

**Figure 4.**
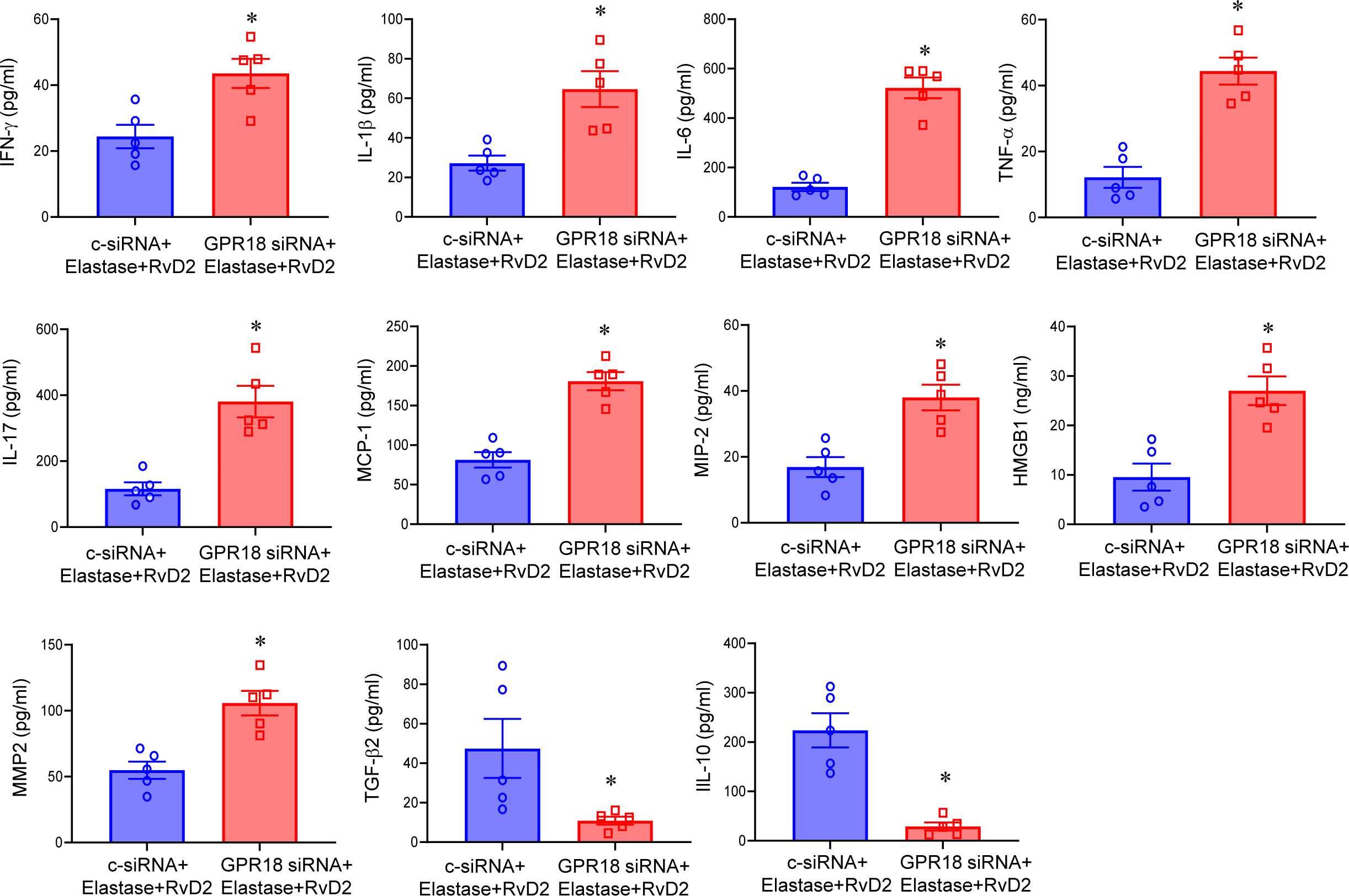
Aortic inflammation is mitigated by RvD2/GPR18 signaling. Aortic tissue from elastase-treated WT mice after c-siRNA+RvD2 administration showed a significant attenuation in pro-inflammatory cytokine/chemokine production and MMP2 expression, and a significant upregulation of TGF-β2 and IL-10 expression, compared to GPR18-siRNA+RvD2-treated mice. *p<0.02; n=5 per group.

### RvD2 treatment induces increase in infiltration and activation of monocytic-MDSCs to mitigate smooth muscle-dependent inflammation and remodeling

Administration of RvD2 in WT mice displayed a significant increase in the M-MDSC population (CD45^+^CD11b^+^CD11c^-^Ly6G^-^Ly6C^+^iNOS^+^) infiltrating in the aortic tissue on day 3 compared to untreated mice (9.5±1.3 vs. 4.1±0.7%; p=0.001; **Figure 5**). No significant differences were observed in M-MDSC quantification on days 7 and 14 in the RvD2-treated mice compared to untreated controls (**Supplemental Figure S3**). These results highlight a critical association between RvD2-mediated protection and M-MDSC infiltration during the inflammation-resolution process. To investigate the mechanistic crosstalk between RvD2-activated M-MDSC and aortic parenchyma, we used *in vitro* studies involving murine M-MDSCs (isolated as shown in **Supplemental Figure S4**) and SMCs. Transient elastase-treatment with/without RvD2 was performed on cultured M-MDSCs that demonstrated a significant increase in TGF-β2 and IL-10 expression in culture supernatants after 24hrs (**Figure 6A-B**). Furthermore, the RvD2-dependent increase in TGF-β2 and IL-10 secretion was abolished when cells were pretreated with GPR18-siRNA compared to M-MDSCs treated with RvD2 and GPR18-siRNA (**Figure 6A-B**). No significant change in expression of TGF-β2 or IL-10 was observed in RvD2-treated cultures of G-MDSCs (**Supplemental Figure S5**).

**Figure 5.**
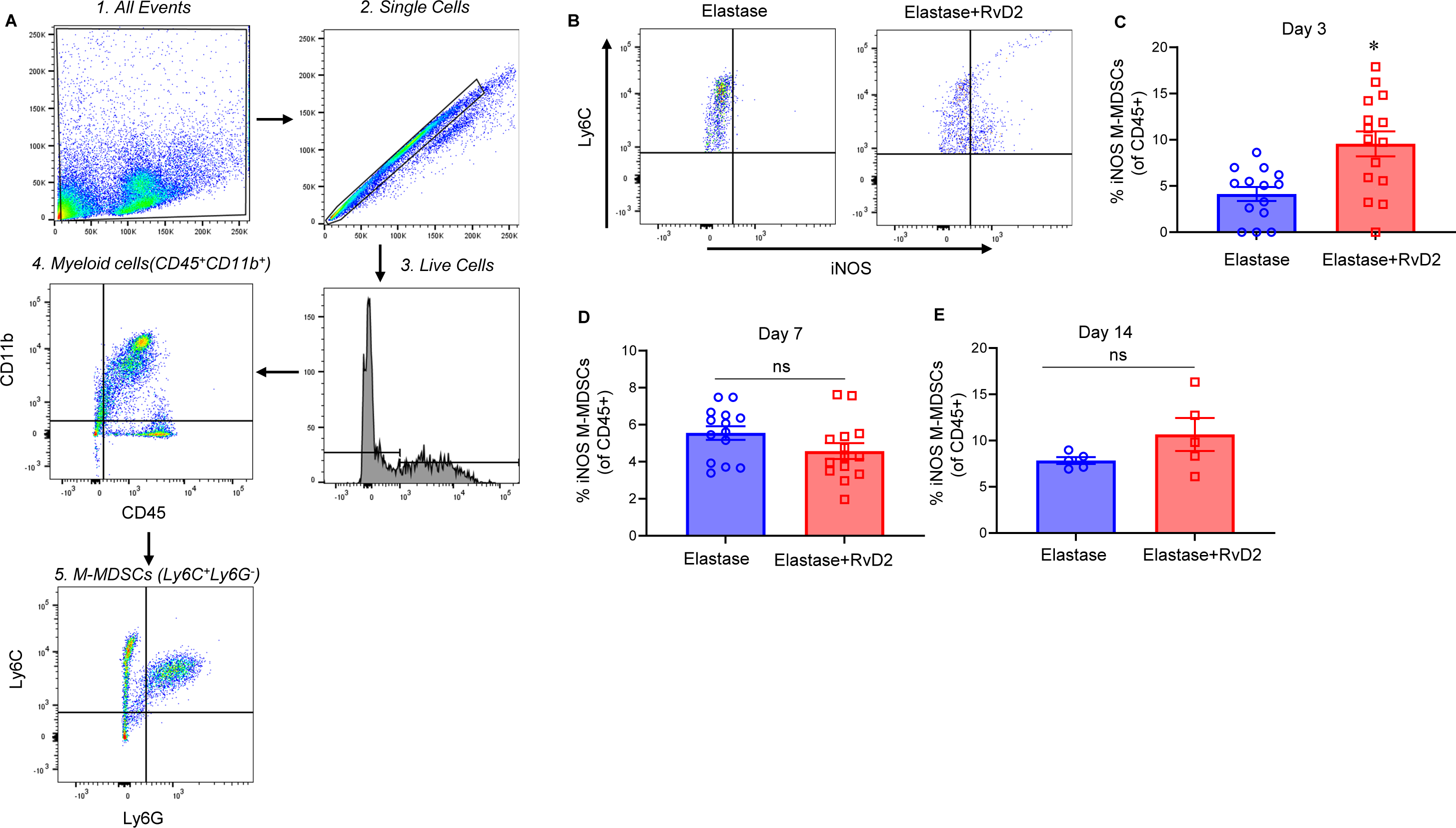
**A**, Representative flow cytometry gating strategy for M-MDSC identification in aortic tissue. **B**, The percentage of M-MDSCs was significantly upregulated in elastase-treated mice administered with RvD2 compared to elastase-treated mice alone on day 3. **C-E**, No significant differences were observed in M-MDSC quantification on days 7 and 14 between these groups. *p=0.001; n=5-15 per group; ns, not significant.

**Figure 6.**
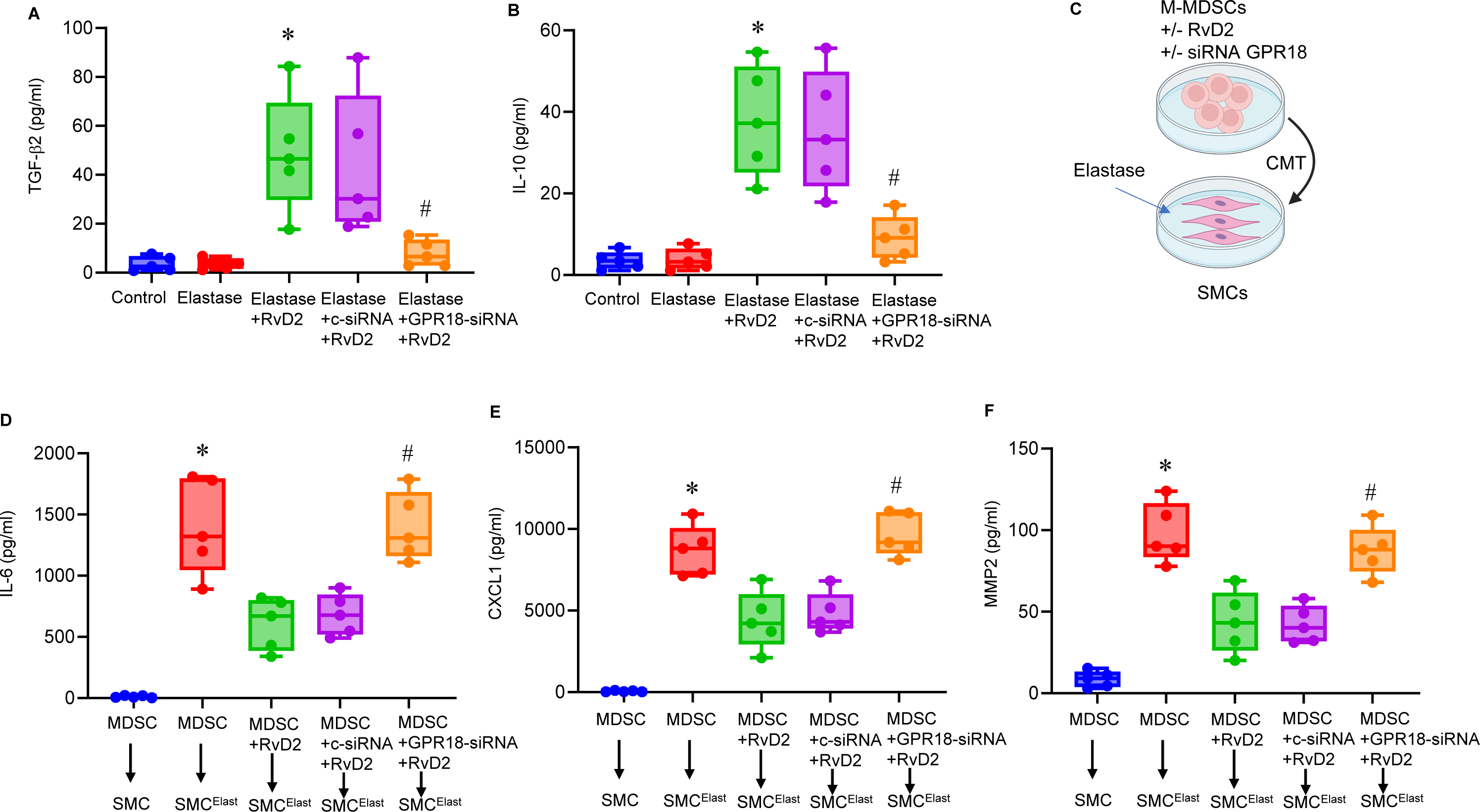
RvD2 upregulates M-MDSC-mediated TGF-β2 and IL-10 secretion to decrease SMC activation. **A-B**, RvD2 treatment upregulates M-MDSC-mediated TGF-β2 and IL-10 secretion in a GPR18-dependent manner compared to untreated cells. *p<0.01 vs control and elastase; #p<0.01 vs. c-siRNA+elastase+RvD2; n=5/group. **C**, Schematic of conditioned media transfer (CMT) experiments between M-MDSCs and SMC cultures. **D-F**, CMT from RvD2-treated M-MDSCs to elastase-treated SMCs significantly mitigated IL-6, CXCL1 and MMP2 expression compared to CMT from untreated MDSCs to elastase-treated SMCs. Inhibition of SMC secreted cytokine and MMP2 expression by RvD2-treated M-MDSCs is dependent on GPR18 receptors. M-MDSCs treated with GPR18-siRNA+RvD2 abolished the decrease in SMC secreted cytokine and MMP2 expression compared to CMT from control (c) siRNA+RvD2. *p<0.001 vs. MDSC+RvD2→SMC^elastase^ and MDSC→SMC; #p<0.001 vs. MDSC+c-siRNA+RvD2→SMC^elastase^; n=5 per group. SMC^elast^ denotes SMC^elastase^.

In a separate group of experiments, conditioned media transfer (CMT) was performed between M-MDSCs and SMCs (**Figure 6C**). M-MDSCs treated with/without RvD2 underwent CMT to SMCs treated with/without transient elastase followed by analysis of cell culture supernatants for cytokines and MMP2 expression. Elastase-treatment induced significant expression of IL-6, CXCL1, and MMP2 in SMCs which was abolished after CMT from RvD2-treated M-MDSCs in a GPR18-dependent manner (**Figure 6D-F**). No change in expression of TGF-β2 and IL-10 expression was observed on CMT from elastase-treated SMCs to M-MDSCs, signifying that the crosstalk between M-MDSCs and SMCs is unidirectional (**Supplemental Figure S6**). Taken together, these results demonstrate the ability of RvD2-mediated immunosuppression by enhancing the infiltration and activation of GPR18 on M-MDSCs to attenuate SMC activation and substantially mitigate aortic inflammation and remodeling during AAA formation (**Figure 7**).

**Figure 7.**
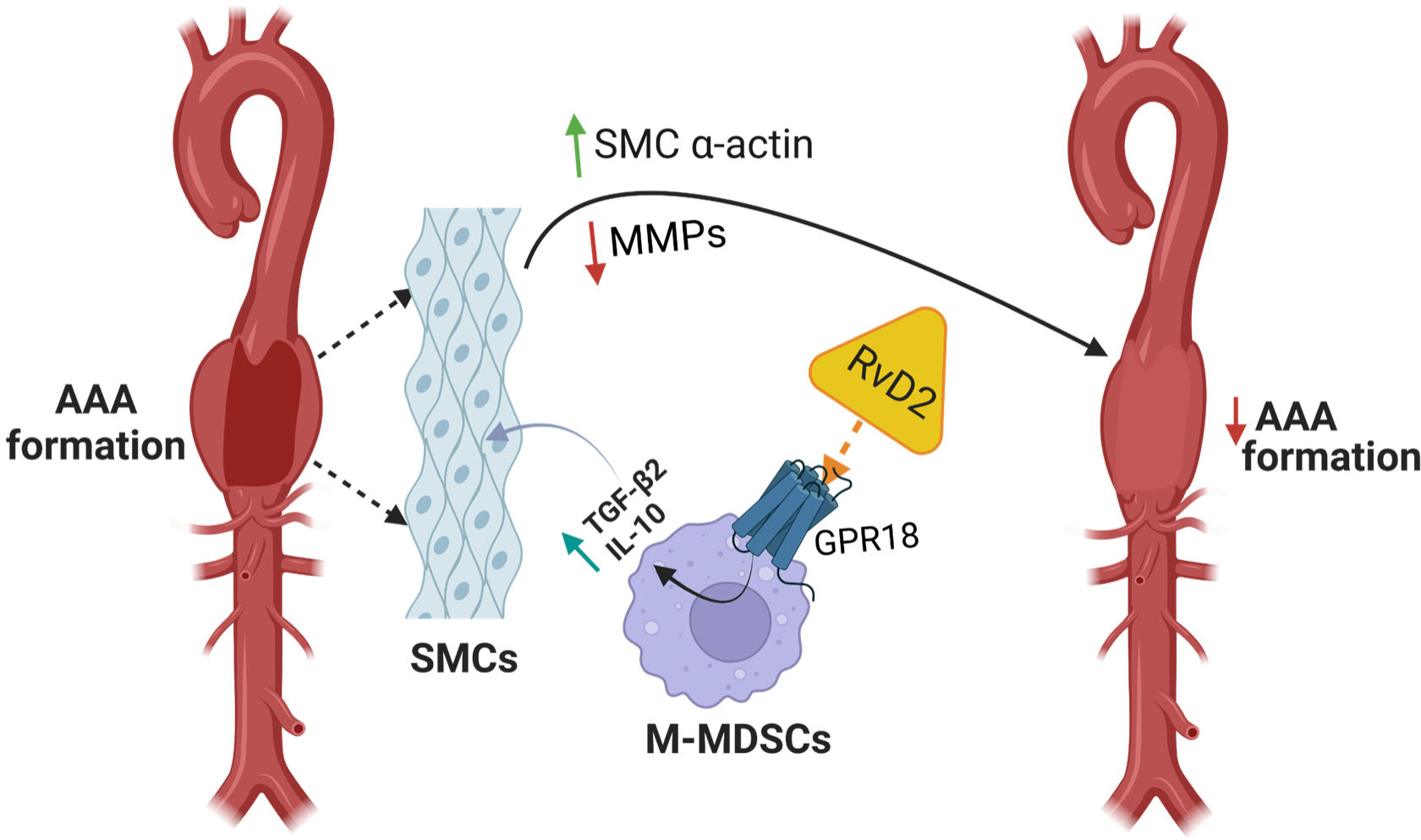
Schematic description of RvD2 mediated attenuation of aortic inflammation and vascular remodeling during AAA formation. RvD1 acts on GPR18 receptors on M-MDSCs to upregulate TGF-β2 and IL-10 secretion leading to inhibition of SMC activation and MMP2 expression. This crosstalk between infiltrating RvD2-induced M-MDSCs and aortic SMCs modulates subsequent aortic remodeling causing attenuation of AAA formation. RvD2, resolvin D2; GPR18, G-protein coupled receptor 18; SMCs; smooth muscle cells; M-MDSCs; monocytic myeloid-derived suppressor cells; MMP; matrix metalloproteinases.

## DISCUSSION

The process of inflammation-resolution plays a pivotal role in chronic vascular pathologies in limiting the disease process in the presence of cytotoxic stimuli. In the context of cardiovascular pathologies, dysregulation of endogenous resolution pathways can lead to uncontrolled aortic tissue inflammation and subsequent remodeling (39–41). In this study, we characterized the role of specialized pro-resolving lipid mediator, RvD2, and the downstream signaling executed via the receptor, GPR18, in upregulating resolution pathways via enhancement of myeloid-precursors cell subsets to attenuate AAA formation. Our results demonstrated that RvD2 treatment can effectively decrease aortic inflammation, leukocyte transmigration, and vascular remodeling during AAA formation in a GPR18-dependent manner. Moreover, we observed a significant increase in myeloid-derived suppressor cells due to RvD2 treatment, especially in the early phase of aortic inflammation. Mechanistically, we further defined that RvD2/GPR18 axis treatment can upregulate monocytic(M)-MDSC activation that leads to increase in TGF-β2 and IL-10 secretion. Finally, the crosstalk between M-MDSCs and aortic SMCs demonstrated the ability of RvD2/GPR18-mediated attenuation of vascular remodeling via inactivation of SMC-specific cytokines and MMP2 expression. Collectively, these results demonstrates the ability of RvD2/GPR18 signaling to result in increased myelopoiesis, enhanced M-MDSC infiltration and activation, thereby promoting the resolution of inflammation during AAA formation.

The understanding of the process of resolution of inflammation (also known as catabasis) has grown significantly in recent years and increasing numbers of key molecules involved have been identified(42). Recent evidence suggests that resolution is an active process with the critical components involving inactivation and cessation of recruitment of neutrophils as well as promotion of macrophage recruitment, efferocytosis, and clearance of pathogens and cellular debris(43) A special class of lipids called specialized pro-resolving mediators (SPMs) are molecules derived from polyunsaturated fatty acids that are produced during the acute inflammatory phase as well as during efferocytosis, and function to resolve inflammation by promoting the core functions of resolution of inflammation(10, 11). SPMs are derived from ω-3 and ω-6 fatty acids and can be organized into five classes of molecules, lipoxins, E-series and D-series resolvins, protectins and maresins(11). These pro-resolving molecules exert their influence in pico-to low nano-molar concentrations via cell surface G protein-coupled receptors including ALX/FPR2, DRV1/GPR32, DRV2/GPR18, ERV1/ChemR23, GPR37, GPR101, and LGR6 and act on various parenchymal and immune cells including neutrophils, monocytes, macrophages, natural killer cells, dendritic cells, epithelial cells, and endothelial cells(10). One of the key mechanisms of SPM-mediated signaling includes facilitating macrophage phagocytosis of cellular debris and inhibition of pro-inflammatory functions via cytokine and chemokine signaling. In order to promote resolution of inflammation, SPMs have the ability to enhance M1/M2 phenotypic polarization in macrophages(13). However, the mechanism that determines the class switch from pro-inflammatory lipid mediators to resolving lipid mediators is not well defined as the process of aging appears to have an impact on SPM biosynthesis, which inhibits or slows the resolution of inflammation.

Based on the current understanding of the role of inflammation in the underlying pathophysiology of vascular pathologies, a few of the bioactive isoforms of SPMs have been previously investigated. In particular, Resolvin D1 and Maresin-1 appear to have the most prominent and well understood role in controlling vascular diseases via downregulation of pro-inflammatory signaling and upregulation of macrophage-dependent efferocytosis(44–46). It has been demonstrated in human and murine models that SPM expressions in plasma or inflamed tissue is associated with disease incidence indicating a correlative association between lipid mediators and vascular pathologies(47, 48). In the context of AAAs, there is evidence that polyunsaturated fatty acid supplementation in animal models reduces inflammation and degeneration of the matrix. Previous studies from our group and others have shown that RvD1 and RvD2 promote an M2 phenotype in macrophages in disease models and reduce matrix metalloproteinases, decreasing the size of aneurysms in mice(13, 45). Additionally, the reparative role of MaR1 has also been shown to significantly attenuate growth of aortic aneurysms by upregulating the process of efferocytosis(34). However, the mechanistic aspects of RvD2-mediated signaling in vascular diseases remains to be deciphered. Recent data suggests that RvD2 may exert bone marrow to exert a pro-resolving function in the liver and endogenously produced within the bone marrow upon inflammation(49). Thus, local production and signaling of SPMs like RvD2 within the bone could be an additional mechanism for the protective actions of RvD2-mediated signaling. Our data in this study also confirms that exogenous RvD2 administration has the ability to enhance bone marrow function by producing monocytic and granulocytic progenitors. The immunosuppression activity of monocytic MDSCs was observed to play a critical role in mediating inhibition of vascular inflammation by acting on SMCs, whereas the granulocytic component of MDSCs failed to do so. These findings are critical to understand the myriad of immunological activities in the biology of myeloid and granulocytic subsets during emergency myelopoiesis that may differ based on disease pathologies, and need to be further investigated.

Monocytic subset of MDSCs are readily recruited to sites of inflammation through canonical trafficking pathways like the CCL2/CCR2 axis (50). These regulatory cells are capable of mediating immune suppression through secretion of anti-inflammatory cytokines, recruitment of regulatory immune cells, exhaustion of pro-inflammatory cells through nutrient sequestering, and facilitating cell polarization to anti-inflammatory phenotypes (20). The functional capabilities of monocytes/macrophages are affected by cleavage of efferocytic receptors, such as TAM (Tyro3, Axl, MerTK), in the inflammatory tissue milieu(51). Therefore, it becomes imperative to discover therapeutic modalities that can enhance endogenous immune regulation. Although previous studies have indicated the role of SPMs like RvD1 in enhancing protective capabilities of macrophages by preserving the efferocytic capabilities, our study pinpoints the previously undescribed role of RvD2-dependent signaling in enhancing endogenous resolution via increase in bone marrow precursors of monocytic compartment that precede the M2 macrophage phenotype and readily immunomodulate the aortic tissue microenvironment. This alternate strategy supplements the phenotype switch of M1/M2 mature macrophages for inducing an anti-inflammatory profile, and provides an endogenous resolution to tissue inflammation by providing a monocytic subset that readily provides an immunocompetent profile, A few limitations of this study should be considered for a translational strategy. The murine elastase AAA model is an excellent experimental tool to delineate the early inflammatory signaling pathways in the aortic wall during AAA, as the important hallmarks of human aneurysm pathology such as macrophage infiltration, matrix degradation, increased MMP activation, elastin fiber degradation, and loss of smooth muscle integrity are recapitulated. However, this experimental model lacks progression of AAA to aortic rupture and chronic vascular inflammation and remodeling as observed clinically (52). Therefore, the findings observed in this study need to be further delineated in chronic aortic aneurysm and rupture models, as we recently described (53, 54). Furthermore, a combined therapeutic strategy using additional bioactive isoforms of SPMs such as MaR1, in combination with RvD2, represents a multi-faceted approach for a combined effective attenuation of aortic inflammation and vascular remodeling to mitigate AAAs (34). Future studies will be focused on using the mechanistic approach of MaR1-dependent upregulation of efferocytosis in combination with RvD2-mediated decrease in SMC activation.

In summary, these findings suggest that the pro-resolving lipid mediator RvD2 plays an important role in enhancing the endogenous inflammation-resolution pathways by enhancing the monocytic compartment of macrophage precursors that mitigate SMC-regulated vascular remodeling and AAA formation. Administration of RvD2-mediated protection is attributable to GPR18-dependent signaling via increase in infiltration and activation of M-MDSCs to mitigate downstream tissue inflammation and remodeling of the aortic tissue. The enhancement of resolution via therapeutic strategies represents an important clinical translational approach as anti-inflammatory treatments can reduce host defense mechanisms leading to enhanced susceptibility of infections that are especially critical in the elderly population. Our results suggest a previously undescribed mechanism to boost endogenous resolution pathways by using bioactive SPMs like RvD2, that can be harnessed to alter the inflammation-resolution dysregulation, for management of chronic inflammatory vascular diseases.

## DATA AVAILABILITY STATEMENT

All relevant data within the manuscript and in Supporting information are available upon request from corresponding authors.

## AUTHOR CONTRIBUTIONS

Ashish K. Sharma and Gilbert R. Upchurch Jr designed the research; Paolo Bellotti, Zachary Ladd, Victoria Leroy, Gang Su, Shiven Sharma, Joseph Hartman, Jonathan Krebs, Chelsea Viscardi, and Ashish K. Sharma performed the research; Paolo Bellotti, Zachary Ladd, Victoria Leroy, Robert Maile, Lyle L. Moldawer, Philip A. Efron, Ashish K. Sharma and Gilbert R. Upchurch Jr analyzed the data; Ashish K. Sharma wrote the manuscript with contributions from all authors; and all authors reviewed and approved the final manuscript.

## ACKNOWLEDGMENTS

This study was supported by NIH R01HL153341 and R01HL138931 (GRU and AKS). Schematic figures were made using Biorender.

## Abbreviations

AAA: abdominal aortic aneurysm
RvD2: resolvin D2
GPR18: formyl peptide receptor
MDSCs: myeloid-derived suppressor cells
α-SMA: alpha-smooth muscle cell actin
HMGB1: high mobility group box 1

## REFERENCES

1. Ailawadi, G., Eliason, J. L., and Upchurch, G. R., Jr. (2003) Current concepts in the pathogenesis of abdominal aortic aneurysm. J Vasc Surg 38, 584–588

2. Bengtsson, H., Sonesson, B., and Bergqvist, D. (1996) Incidence and prevalence of abdominal aortic aneurysms, estimated by necropsy studies and population screening by ultrasound. Ann N Y Acad Sci 800, 1–24

3. Dimick, J. B., Stanley, J. C., Axelrod, D. A., Kazmers, A., Henke, P. K., Jacobs, L. A., Wakefield, T. W., Greenfield, L. J., and Upchurch, G. R., Jr. (2002) Variation in death rate after abdominal aortic aneurysmectomy in the United States: impact of hospital volume, gender, and age. Ann Surg 235, 579–585

4. Johnston, K. W., Rutherford, R. B., Tilson, M. D., Shah, D. M., Hollier, L., and Stanley, J. C. (1991) Suggested standards for reporting on arterial aneurysms. Subcommittee on Reporting Standards for Arterial Aneurysms, Ad Hoc Committee on Reporting Standards, Society for Vascular Surgery and North American Chapter, International Society for Cardiovascular Surgery. J Vasc Surg 13, 452–458

5. Anidjar, S., Dobrin, P. B., Eichorst, M., Graham, G. P., and Chejfec, G. (1992) Correlation of inflammatory infiltrate with the enlargement of experimental aortic aneurysms. J Vasc Surg 16, 139–147

6. Aziz, F., and Kuivaniemi, H. (2007) Role of matrix metalloproteinase inhibitors in preventing abdominal aortic aneurysm. Ann Vasc Surg 21, 392–401

7. Freestone, T., Turner, R. J., Coady, A., Higman, D. J., Greenhalgh, R. M., and Powell, J. T. (1995) Inflammation and matrix metalloproteinases in the enlarging abdominal aortic aneurysm. Arterioscler Thromb Vasc Biol 15, 1145–1151

8. Crowther, M., Goodall, S., Jones, J. L., Bell, P. R., and Thompson, M. M. (2000) Increased matrix metalloproteinase 2 expression in vascular smooth muscle cells cultured from abdominal aortic aneurysms. J Vasc Surg 32, 575–583

9. Xiong, W., Knispel, R., Mactaggart, J., and Baxter, B. T. (2006) Effects of tissue inhibitor of metalloproteinase 2 deficiency on aneurysm formation. J Vasc Surg 44, 1061–1066

10. Serhan, C. N. (2014) Pro-resolving lipid mediators are leads for resolution physiology. Nature 510, 92–101

11. Basil, M. C., and Levy, B. D. (2016) Specialized pro-resolving mediators: endogenous regulators of infection and inflammation. Nat Rev Immunol 16, 51–67

12. Chiang, N., de la Rosa, X., Libreros, S., and Serhan, C. N. (2017) Novel Resolvin D2 Receptor Axis in Infectious Inflammation. J Immunol 198, 842–851

13. Pope, N. H., Salmon, M., Davis, J. P., Chatterjee, A., Su, G., Conte, M. S., Ailawadi, G., and Upchurch, G. R., Jr. (2016) D-series resolvins inhibit murine abdominal aortic aneurysm formation and increase M2 macrophage polarization. FASEB J 30, 4192–4201

14. Zhang, J., Hodges, A., Chen, S. H., and Pan, P. Y. (2021) Myeloid-derived suppressor cells as cellular immunotherapy in transplantation and autoimmune diseases. Cell Immunol 362, 104300

15. Scalea, J. R., Lee, Y. S., Davila, E., and Bromberg, J. S. (2018) Myeloid-Derived Suppressor Cells and Their Potential Application in Transplantation. Transplantation 102, 359–367

16. Zhang, W., Li, J., Qi, G., Tu, G., Yang, C., and Xu, M. (2018) Myeloid-derived suppressor cells in transplantation: the dawn of cell therapy. J Transl Med 16

17. Yaseen, M. M., Abuharfeil, N. M., Darmani, H., and Daoud, A. (2020) Mechanisms of immune suppression by myeloid-derived suppressor cells: the role of interleukin-10 as a key immunoregulatory cytokine. Open Biol 10, 200111

18. Sinha, P., Clements, V. K., Bunt, S. K., Albelda, S. M., and Ostrand-Rosenberg, S. (2007) Cross-talk between myeloid-derived suppressor cells and macrophages subverts tumor immunity toward a type 2 response. J Immunol 179, 977–983

19. Veglia, F., Perego, M., and Gabrilovich, D. (2018) Myeloid-derived suppressor cells coming of age. Nat Immunol 19, 108–119

20. Gabrilovich, D. I., and Nagaraj, S. (2009) Myeloid-derived suppressor cells as regulators of the immune system. Nat. Rev. Immunol. 9, 162–174

21. Mazzoni, A., Bronte, V., Visintin, A., Spitzer, J. H., Apolloni, E., Serafini, P., Zanovello, P., and Segal, D. M. (2002) Myeloid Suppressor Lines Inhibit T Cell Responses by an NO-Dependent Mechanism. The Journal of Immunology 168, 689

22. Bronte, V., Serafini, P., Mazzoni, A., Segal, D. M., and Zanovello, P. (2003) L-arginine metabolism in myeloid cells controls T-lymphocyte functions. Trends Immunol 24, 302–306

23. Gabrilovich, D. I., and Nagaraj, S. (2009) Myeloid-derived suppressor cells as regulators of the immune system. Nat Rev Immunol 9, 162–174

24. Nagaraj, S., Schrum, A. G., Cho, H. I., Celis, E., and Gabrilovich, D. I. (2010) Mechanism of T cell tolerance induced by myeloid-derived suppressor cells. J Immunol 184, 3106–3116

25. Srivastava, M. K., Sinha, P., Clements, V. K., Rodriguez, P., and Ostrand-Rosenberg, S. (2010) Myeloid-derived suppressor cells inhibit T-cell activation by depleting cystine and cysteine. Cancer Res 70, 68–77

26. Nagaraj, S., Youn, J.-I., and Gabrilovich, D. I. (2013) Reciprocal Relationship between Myeloid-Derived Suppressor Cells and T Cells. The Journal of Immunology 191, 17–23

27. Schrijver, I. T., Théroude, C., and Roger, T. (2019) Myeloid-Derived Suppressor Cells in Sepsis. Front Immunol 10, 327–327

28. Dysthe, M., and Parihar, R. (2020) Myeloid-Derived Suppressor Cells in the Tumor Microenvironment. Adv Exp Med Biol 1224, 117–140

29. Melero-Jerez, C., Ortega, M. C., Moliné-Velázquez, V., and Clemente, D. (2016) Myeloid derived suppressor cells in inflammatory conditions of the central nervous system. Biochimica et Biophysica Acta (BBA) - Molecular Basis of Disease 1862, 368–380

30. Yi, H., Guo, C., Yu, X., Zuo, D., and Wang, X. Y. (2012) Mouse CD11b+Gr-1+ myeloid cells can promote Th17 cell differentiation and experimental autoimmune encephalomyelitis. J Immunol 189, 4295–4304

31. Ostrand-Rosenberg, S., Sinha, P., Beury, D. W., and Clements, V. K. (2012) Cross-talk between myeloid-derived suppressor cells (MDSC), macrophages, and dendritic cells enhances tumor-induced immune suppression. Semin Cancer Biol 22, 275–281

32. Filiberto, A. C., Spinosa, M. D., Elder, C. T., Su, G., Leroy, V., Ladd, Z., Lu, G., Mehaffey, J. H., Salmon, M. D., Hawkins, R. B., Ravichandran, K. S., Isakson, B. E., Upchurch, G. R., Jr., and Sharma, A. K. (2022) Endothelial pannexin-1 channels modulate macrophage and smooth muscle cell activation in abdominal aortic aneurysm formation. Nat Commun 13, 1521

33. Filiberto, A. C., Ladd, Z., Leroy, V., Su, G., Elder, C. T., Pruitt, E. Y., Hensley, S. E., Lu, G., Hartman, J. B., Zarrinpar, A., Sharma, A. K., and Upchurch, G. R., Jr. (2022) Resolution of inflammation via RvD1/FPR2 signaling mitigates Nox2 activation and ferroptosis of macrophages in experimental abdominal aortic aneurysms. FASEB J 36, e22579

34. Elder, C. T., Filiberto, A. C., Su, G., Ladd, Z., Leroy, V., Pruitt, E. Y., Lu, G., Jiang, Z., Sharma, A. K., and Upchurch, G. R., Jr. (2021) Maresin 1 activates LGR6 signaling to inhibit smooth muscle cell activation and attenuate murine abdominal aortic aneurysm formation. FASEB J 35, e21780

35. Bronte, V., Brandau, S., Chen, S.-H., Colombo, M. P., Frey, A. B., Greten, T. F., Mandruzzato, S., Murray, P. J., Ochoa, A., Ostrand-Rosenberg, S., Rodriguez, P. C., Sica, A., Umansky, V., Vonderheide, R. H., and Gabrilovich, D. I. (2016) Recommendations for myeloid-derived suppressor cell nomenclature and characterization standards. Nature Communications 7, 12150

36. Eckert, I., Ribechini, E., and Lutz, M. B. (2021) In Vitro Generation of Murine Myeloid-Derived Suppressor Cells, Analysis of Markers, Developmental Commitment, and Function. In Myeloid-Derived Suppressor Cells (Brandau, S., and Dorhoi, A., eds) pp. 99–114, Springer US, New York, NY

37. Leroy, V., Manual Kollareth, D. J., Tu, Z., Valisno, J. A. C., Woolet-Stockton, M., Saha, B., Emtiazjoo, A. M., Rackauskas, M., Moldawer, L. L., Efron, P. A., Cai, G., Atkinson, C., Upchurch, G. R., and Sharma, A. K. (2024) MerTK-dependent efferocytosis by monocytic-MDSCs mediates resolution of post-lung transplant injury. bioRxiv

38. Eckert, I., Ribechini, E., and Lutz, M. B. (2021) In Vitro Generation of Murine Myeloid-Derived Suppressor Cells, Analysis of Markers, Developmental Commitment, and Function. Methods Mol Biol 2236, 99–114

39. Maskrey, B. H., Megson, I. L., Whitfield, P. D., and Rossi, A. G. (2011) Mechanisms of resolution of inflammation: a focus on cardiovascular disease. Arterioscler Thromb Vasc Biol 31, 1001–1006

40. Fredman, G. (2019) Can Inflammation-Resolution Provide Clues to Treat Patients According to Their Plaque Phenotype? Front Pharmacol 10, 205

41. Halade, G. V., and Lee, D. H. (2022) Inflammation and resolution signaling in cardiac repair and heart failure. EBioMedicine 79, 103992

42. Widgerow, A. D. (2012) Cellular resolution of inflammation--catabasis. Wound Repair Regen 20, 2–7

43. Panigrahy, D., Gilligan, M. M., Serhan, C. N., and Kashfi, K. (2021) Resolution of inflammation: An organizing principle in biology and medicine. Pharmacol Ther 227, 107879

44. Olivares-Silva, F., De Gregorio, N., Espitia-Corredor, J., Espinoza, C., Vivar, R., Silva, D., Osorio, J. M., Lavandero, S., Peiro, C., Sanchez-Ferrer, C., and Diaz-Araya, G. (2021) Resolvin-D1 attenuation of angiotensin II-induced cardiac inflammation in mice is associated with prevention of cardiac remodeling and hypertension. Biochim Biophys Acta Mol Basis Dis 1867, 166241

45. Spinosa, M., Su, G., Salmon, M. D., Lu, G., Cullen, J. M., Fashandi, A. Z., Hawkins, R. B., Montgomery, W., Meher, A. K., Conte, M. S., Sharma, A. K., Ailawadi, G., and Upchurch, G. R., Jr. (2018) Resolvin D1 decreases abdominal aortic aneurysm formation by inhibiting NETosis in a mouse model. J Vasc Surg 68, 93S–103S

46. Viola, J. R., Lemnitzer, P., Jansen, Y., Csaba, G., Winter, C., Neideck, C., Silvestre-Roig, C., Dittmar, G., Doring, Y., Drechsler, M., Weber, C., Zimmer, R., Cenac, N., and Soehnlein, O. (2016) Resolving Lipid Mediators Maresin 1 and Resolvin D2 Prevent Atheroprogression in Mice. Circ Res 119, 1030–1038

47. Salazar, J., Pirela, D., Nava, M., Castro, A., Angarita, L., Parra, H., Duran-Aguero, S., Rojas-Gomez, D. M., Galban, N., Anez, R., Chacin, M., Diaz, A., Villasmil, N., De Sanctis, J. B., and Bermudez, V. (2022) Specialized Proresolving Lipid Mediators: A Potential Therapeutic Target for Atherosclerosis. Int J Mol Sci 23

48. Kim, A. S., and Conte, M. S. (2020) Specialized pro-resolving lipid mediators in cardiovascular disease, diagnosis, and therapy. Adv Drug Deliv Rev 159, 170–179

49. Fitzgerald, H., Bonin, J. L., Sadhu, S., Lipscomb, M., Biswas, N., Decker, C., Nabage, M., Bossardi, R., Marinello, M., Mena, A. H., Gilliard, K., Spite, M., Adam, A., MacNamara, K. C., and Fredman, G. (2023) The Resolvin D2-GPR18 Axis Enhances Bone Marrow Function and Limits Hepatic Fibrosis in Aging. bioRxiv

50. Serbina, N. V., and Pamer, E. G. (2006) Monocyte emigration from bone marrow during bacterial infection requires signals mediated by chemokine receptor CCR2. Nat Immunol 7, 311–317

51. Vago, J. P., Amaral, F. A., and van de Loo, F. A. J. (2021) Resolving inflammation by TAM receptor activation. Pharmacol Ther 227, 107893

52. Bhamidipati, C. M., Mehta, G. S., Lu, G., Moehle, C. W., Barbery, C., DiMusto, P. D., Laser, A., Kron, I. L., Upchurch, G. R., Jr., and Ailawadi, G. (2012) Development of a novel murine model of aortic aneurysms using peri-adventitial elastase. Surgery 152, 238–246

53. Fashandi, A. Z., Hawkins, R. B., Salmon, M. D., Spinosa, M. D., Montgomery, W. G., Cullen, J. M., Lu, G., Su, G., Ailawadi, G., and Upchurch, G. R., Jr. (2018) A novel reproducible model of aortic aneurysm rupture. Surgery 163, 397–403

54. Lu, G., Su, G., Davis, J. P., Schaheen, B., Downs, E., Roy, R. J., Ailawadi, G., and Upchurch, G. R., Jr. (2017) A novel chronic advanced stage abdominal aortic aneurysm murine model. J Vasc Surg 66, 232–242 e234

