## Supplemental Figures for "Resolvin D2/GPR18 signaling enhances monocytic myeloid-derived suppressor cell function to mitigate abdominal aortic aneurysm formation"

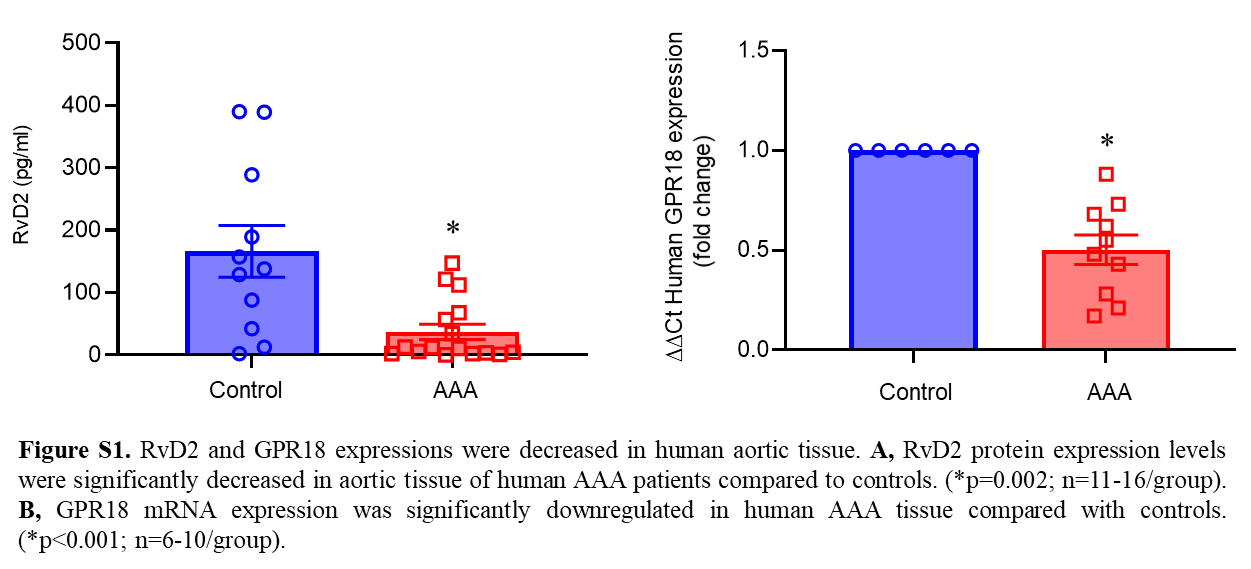
**Supplemental Figure S1**

**Supplemental Figure S2**

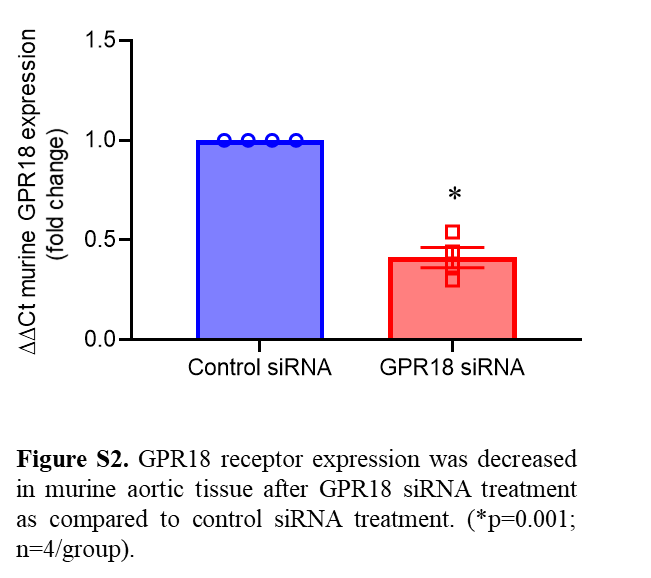

**Supplemental Figure S3**
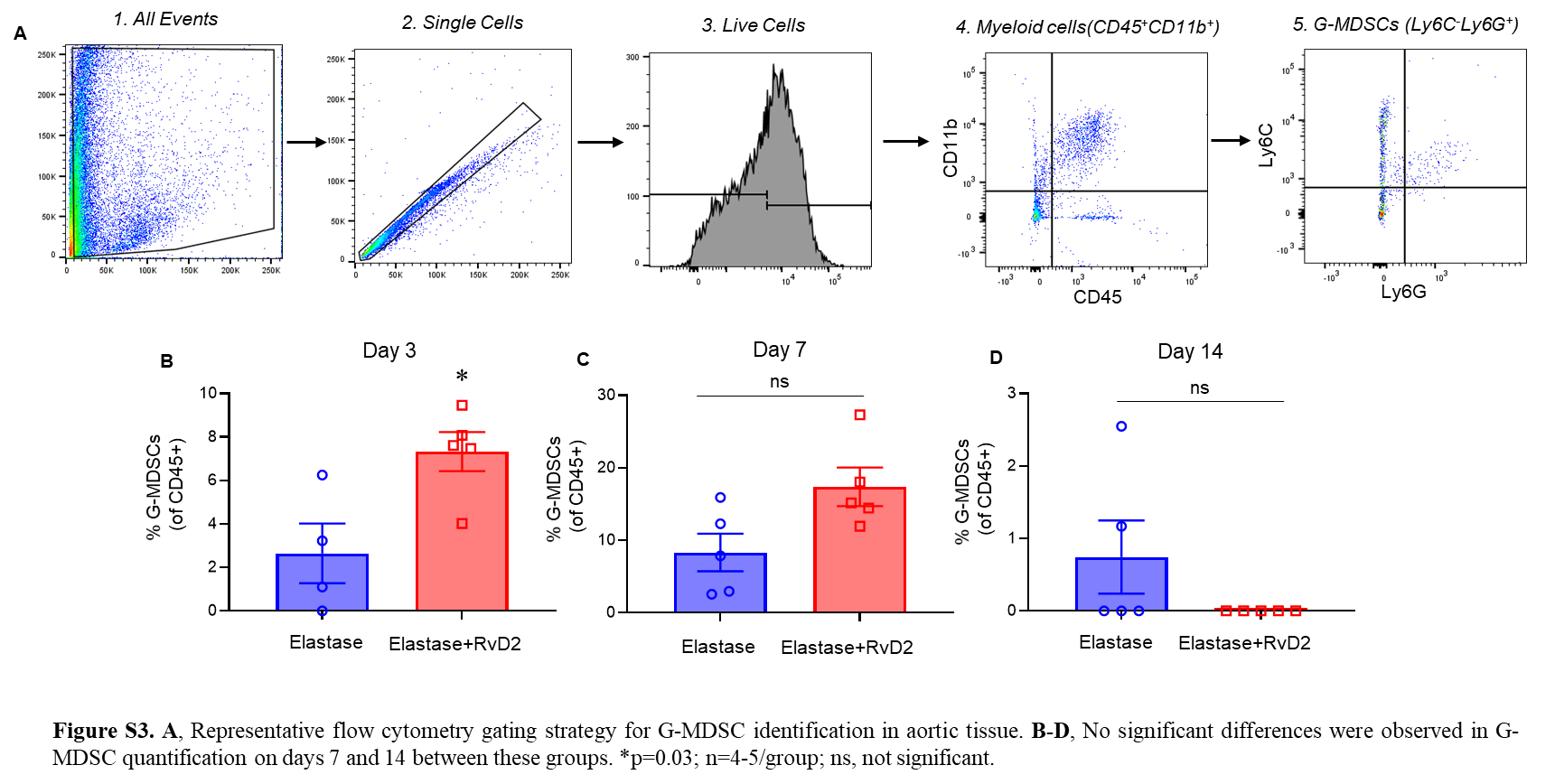

**Supplemental Figure S4**

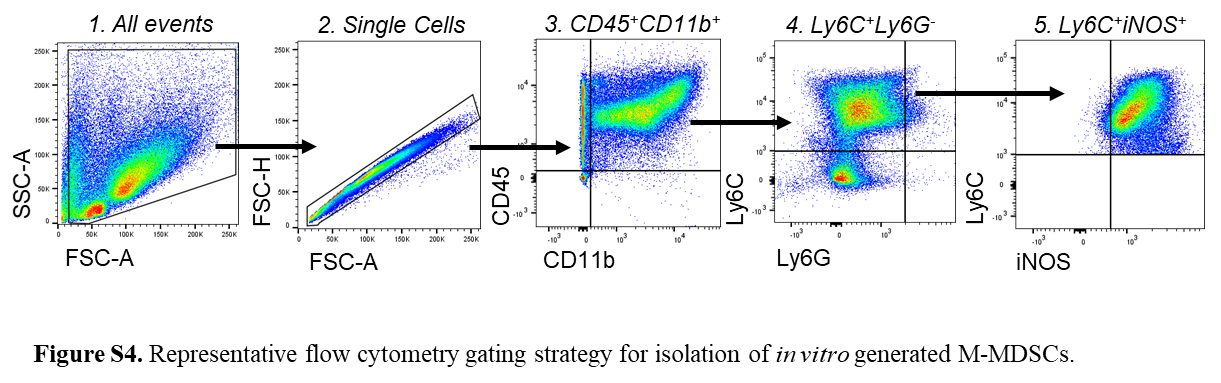

**Supplemental Figure S5**

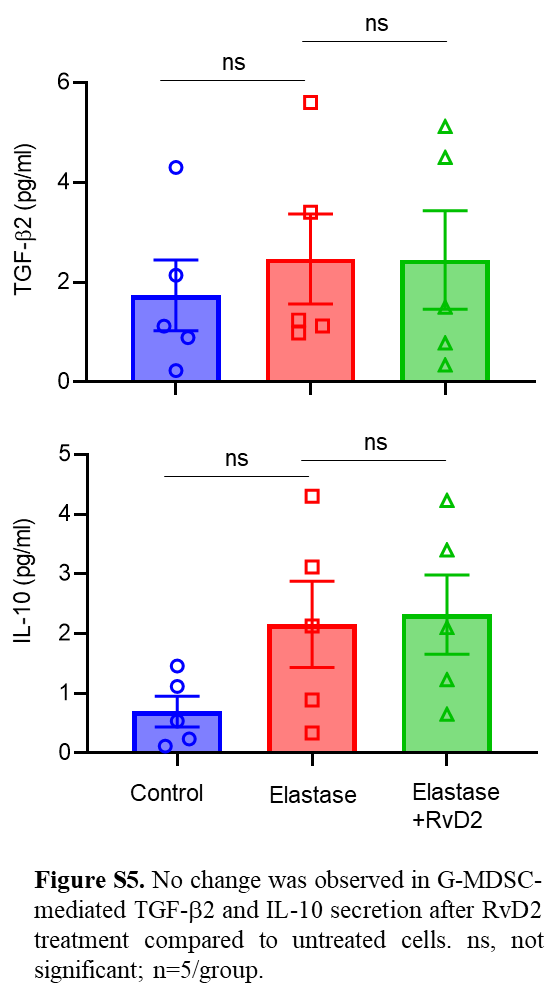

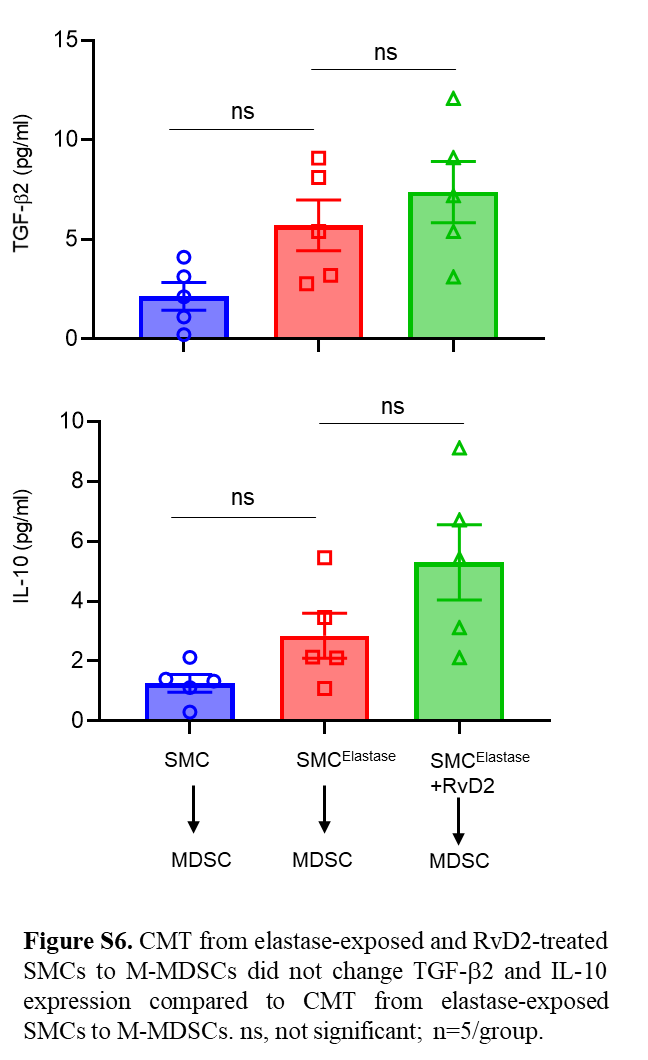
**Supplemental Figure S6**
